# Isolation and expansion of pure and functional γδ T cells

**DOI:** 10.1101/2023.11.12.566762

**Authors:** T. Verkerk, A. T. Pappot, T. Jorritsma, L.A. King, R. M. Spaapen, S. M. van Ham

## Abstract

γδ T cells are important components of the immune system due to their ability to elicit a fast and strong response against infected and transformed cells. Because they can specifically and effectively kill target cells in an MHC independent fashion, there is great interest to utilize these cells in anti-tumor therapies where antigen presentation may be hampered. Since only a small fraction of T cells in the blood or tumor tissue are γδ T cells, they require extensive expansion to allow for fundamental, preclinical and *ex vivo* research. Although expansion protocols can be successful, most are based on depletion of other cell types rather than γδ T cell specific isolation, resulting in unpredictable purity of the isolated fraction. Moreover, the primary focus only lies with expansion of Vδ2^+^ T cells, while Vδ1^+^ T cells likewise have anti-tumor potential. Here, we investigated whether γδ T cells directly isolated from blood could be efficiently expanded while maintaining function. γδ T cell subsets were isolated using MACS separation, followed by FACS sorting, yielding >99% pure γδ T cells. Isolated Vδ1^+^ and Vδ2^+^ T cells could effectively expand immediately after isolation or upon freeze/thawing and reached expansion ratios between 200 to 2000-fold starting from varying numbers using cytokine supported feeder stimulations. After expansion, potential effector functions of γδ T cells were demonstrated by IFN-γ, TNF-α and granzyme B production upon PMA/ionomycin stimulation and effective killing capacity of multiple tumor cell lines was confirmed in killing assays.

In conclusion, pure γδ T cells can productively be expanded while maintaining their anti-tumor effector functions against tumor cells. Moreover, γδ T cells could be expanded from low starting numbers suggesting that this protocol may even allow for expansion of cells extracted from tumor biopsies.

## Introduction

γδ T cells are considered to be potent cells to combat infections and malignancies because of their strategic localization in mucosal tissues, specific recognition of target cells and direct response upon activation. Like αβ T cells, γδ T cells can be activated through their TCR and have the strong capacity to elicit a cytotoxic response composed by release of TNF-α, IFN-γ, perforins and granzymes (*1*–*3*). Unlike αβ TCRs, γδ-TCR triggering is typically independent of antigens presented on major histocompatibility complex I or II (MHC-I/II). In addition, γδ T cells are fully mature when they leave the thymus, have a broad range of antigen recognition and can consequently respond rapidly to antigen encounter. Similar to both αβ T cells and natural killer (NK) cells, γδ T cells express NKG2D which can detect stress-induced signals on tumors and infected cells (*4*). In addition, these cells also share other receptors with the innate NK cells such as FcγRIIIa (CD16) which is required to recognize opsonized tumor cells, DNAM-1 (CD226) and natural cytotoxicity receptors (NCRs) NKp46, NKp44 and NKp30 which promote anti-tumor cytotoxicity (*5, 6*).

γδ T cells can be divided into different subsets based on the composition of the TCR chains of which Vδ1 and Vδ2 are two major γδ T cells subsets found in humans. Both types can be found in peripheral blood of which the majority are Vδ2 T cells. Vδ1 T cells mainly reside in tissues and mucosa, such as the skin, spleen, gut, and lungs (*7*–*10*).

The majority of γδ T cells is activated in an MHC-I independent manner and only a few γδ T cell clones have been reported to become activated by classical MHC-I molecules (*11*–*13*). Vδ1 T cells recognize MICA/B and ULBPs on stressed cells through γδ TCR and NKG2D engagement or CD1c/d on antigen presenting cells (*1, 14*). Vδ2 T cells harbor a TCR that can be activated through surface expression of the butyrophilin (BTN) BTN3A1/BTN2A1 complex, modulated by pathogenic changes in cells (*15, 16*).

The mode of action of γδ T cells in an anti-tumor response comprises direct cytolytic activity via NKG2D, FcyRIII, Fas/Fas-ligand and TNF related apoptosis inducing ligand (TRAIL) pathway resulting in production of perforins, granzymes, IFN-γ and TNF-α (*4, 5, 17*–*20*). In concurrence, γδ T cells can also stimulate other immune cells (*21, 22*), for instance activate NK cells via co-stimulation, promote dendritic cell (DC) maturation and support initiation of a humoral immune response (*23*–*28*).

Interestingly, Vδ1 and Vδ2 subsets can represent up to 50% of CD3^+^ tumor infiltrating lymphocytes (TILs), although this varies between tumor type and stage (*1, 29*–*33*). Recently, Knight and colleagues found increased frequencies of both Vδ1 and Vδ2 T cells in patients with chronic myeloid leukemia (CML) compared to age-matched healthy donors and demonstrated primary CML specific lysis by autologous Vδ1 T cells (*32*). Some studies found correlations between tumor infiltrating γδ T cells with a favorable clinical outcome. It remains to be established though if tumor infiltrating γδ T cells can be used as a prognostic factor (*1, 22, 30, 34, 35*). Motivation for the application of γδ T cells in immunotherapy is driven by the distinctive MHC independent activation, meaning that its application is not restricted to an autologous setting. In addition, they have a potential to be used for tumors showing immune escape through downmodulation of MHC expression. Immunotherapy based on γδ T cells involves *in vivo* stimulation of Vδ2 T cells through intravenous administration of pamidronate or zoledronate, which are bisphosphonates stimulating BTN3A1/BTN2A1 expression, and autologous or allogenic reinfusion of *ex vivo* zoledronate driven expanded γδ T cells (*3, 36*–*38*). These treatments have been shown to have minor adverse effects and to be effective for a number of patients suffering from for example, pancreatic, lung, liver and hematological cancers, especially in combination with other conventional therapies. (*3, 36*).

The low number of γδ T cells in peripheral blood (0.5-5% of T cells) and, especially, in tumor tissue represents a major challenge for their application in fundamental, preclinical and *ex vivo* studies (*39*–*41*). Therefore, expansion of γδ T cells is often applied to study these cells.

Scientific reports in which expanded γδ T cells are used are mainly subset specific and vary in isolation and expansion strategies. Most used methods are based on the use of bisphosphonates zoledronate, pamidronate or bromohydrin pyrophosphate (BrHPP) to activate Vδ2 T cells specifically residing in total PBMCs (*42*–*45*). These compounds inhibit the IPP processing enzyme farnesyl diphosphate synthase (FDPS), consequently increasing IPP levels and subsequent BTN2A1/BTN3A1 surface expression (*15*). Zoledronate based expansion of Vδ2 T cells in total PBMCs can generate over 90% pure Vδ2 T cells, but shows strong variation in yields between donors (*45*–*49*). Other methods focus on TCR activation with, for instance, lectins such as phytohaemagglutinin (PHA) or anti-CD3 (OKT3) with or without anti-CD28 (*50*). Lastly, K562 cells are used as feeder cells which are modified to express various molecules such as CD80, CD86, CD137L and artificial membrane bound cytokines like IL-15 (*44, 51*). Generally, IL-2 is used to promote activation, growth and proliferation, often combined with other cytokines like IL-4, IL-7, IL-15, IL-18 and IL-21 (*44*). Isolation of γδ T cells usually incorporates an αβ T cell depletion step, occasionally combined with an additional NK cells depletion step, either before or after the expansion period.

The issues that accompany the current described methods is that they are variable, solely applicable to specific subsets and may yield unpredictable or impracticable levels purity of the (final) product. Hence, there is a need for a practical protocol that can be used to specifically isolate Vδ1 and Vδ2 T cells followed by an expansion method that assures purity and maintains their functional properties. Standardized isolation and expansion protocols would allow for comparison between studies and benefit the growth of knowledge on γδ T cell differentiation, function and targets.

Here, we report an isolation and expansion method which reproducibly generates high and >99% pure Vδ1 and Vδ2 T cell numbers after 14 days of expansion using PBMC as starting material. This method focuses first on establishing γδ T cell populations through Vδ1 and Vδ2 T cell specific isolation, which can subsequently be expanded to over a 1000-fold increase. In addition, we show that these Vδ2 T cells can effectively be activated and kill tumor cells. Moreover, this method can be used to expand cells from low starting numbers or frozen, previously expanded cells. Together, this allows investigators to reproducibly generate large batches of γδ T cells to study their biology.

## Materials and Methods

### Isolation of γδ T cells from PBMCs

Buffy coats were collected from healthy donors (Sanquin Blood Supply, Amsterdam, the Netherlands) who provided written informed consent for the use of their donation for research. Peripheral blood mononucleated cells (PBMCs) were isolated from buffy coats using Lymphoprep (Axis-Shield PoC AS, Dundee DD2 1XA, Scotland) density gradient. For direct targeted isolation of γδ T cells, PBMCs were incubated with PE conjugated mouse anti-human Vδ1 TCR or APC/PE conjugated mouse anti-human Vδ2 TCR for 30 minutes on ice (*table 1*). The PBMCs were washed with PBS/0.1%BSA and incubated with anti-mouse IgG microbeads (Miltenyi) prior to positive MACS isolation according to manufacturer’s protocol. Subsequently, the collected γδ T cells were purified using a FACS sort for PE^+^ (Vδ1) or PE/APC^+^ (Vδ2) cells. For untouched, pan γδ T cell isolation, the TCRγ/δ+ T Cell Isolation Kit (Miltenyi) was used according to manufacturer’s protocol. The purity of the isolated γδ T cells was assed prior to cell culture.

**Table 1.**
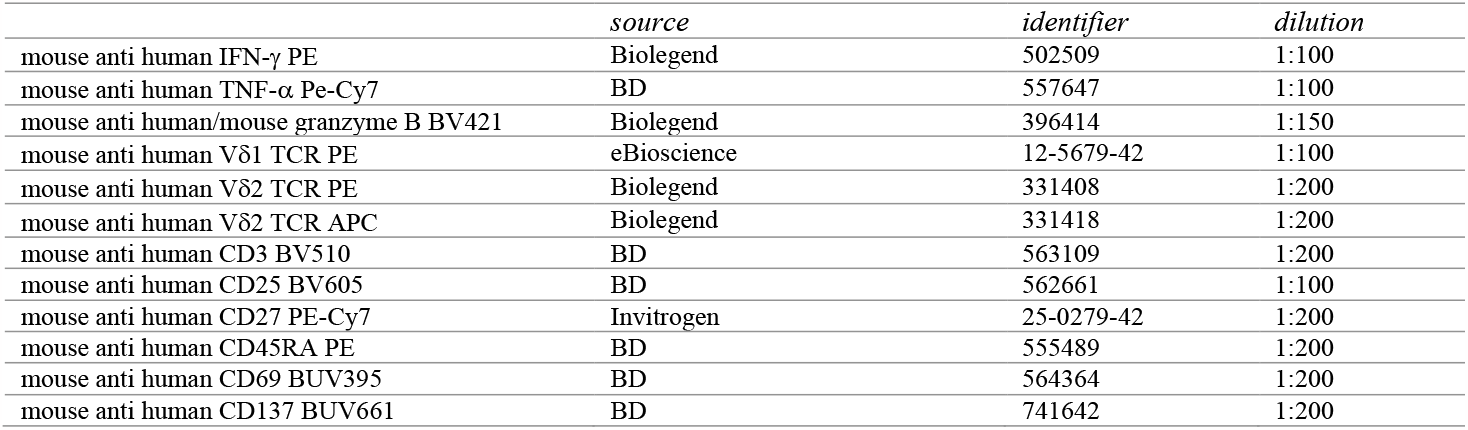
antibodies

### Cell culture

The target cell lines WM9 and WiDr (kindly provided by prof. J.J. van der Vliet, Amsterdam UMC, the Netherlands) were cultured in DMEM (Gibco) supplemented with 10% FCS (Serana), antibiotics (PenStrep, Invitrogen), 1% L-glutamine (Gibco) and 0.05 mM BME (Sigma). The target cell line HAP1 and the EBV-LCLs used as feeder cells were cultured in IMDM (Gibco) supplemented with 10% FCS and antibiotics. Freshly isolated, thawed or 14-day cultured γδ T cells were expanded using the protocol as described in this paper. In short, γδ T cells per well were expanded in IMDM containing feeder cells (0.1×10^6^ irradiated EBV-LCLs (50Gy) and 1×10^6^ PBMCs (30Gy) per ml), 5% heat inactivated human serum (HS, Sanquin), 5% FCS and antibiotics supplemented with 120 U/mL IL-2 (Peprotech), 20 ng/mL IL-7 (Research grade, Miltenyi), 20 ng/mL IL-15 (Peprotech) and 1.0 μg/mL Phytohaemagglutinin (PHA, HA16 Remel™, Thermo Fisher Scientific) at the start of the culture. Cultures with 150 cells started in a 96-well plate in a total volume of 100 μl, 15.000 cells started in a 48-well plate with a total volume of 500 μl and cultures starting at 150.000 cells were performed in 24-well plates with a total volume of 1ml. Cell cultures were harvested, counted and replated in 24-well plates at day 7 and day 10 (15.000 and 150.000 start) or day 17 (150 start) to maintain these cytokine concentrations and a cell density of 0.5 x 10^6^ cells per ml. All cells were cultured at 37°C and 5% CO_2_.

### PMA/Ionomycin stimulation

γδ T cells were stimulated with 100 ng/mL PMA (Sigma-Aldrich) and 1 μg/mL ionomycin (from Streptomyces conglobatus, Sigma-Aldrich) for 1.5 hours at 37 °C and 5% CO_2_. Brefeldin A (1:1000, BD) was added to the medium at the start of stimulation for intracellular measurement of IFN-γ, TNF-α and granzyme B. For analysis, the cells were transferred to a V-bottom plate and washed with cold PBS/0.5%BSA prior to staining. Production of IFN-γ, TNF-α and granzyme B was measured using flowcytometry.

### Coculture assays

The target cells WM9, WiDr and HAP1, were plated one day prior to the coculture with γδ T cells at a density of 15.000 cells/well in a flat-bottom 96-well plate and treated O/N with 10 μM pamidronate (PAM, Sigma) in complete IMDM or DMEM. After the O/N culture, the wells were carefully washed to remove dead cells and freshly expanded γδ T cells were added at a 5:1 ratio (E:T) in a total volume of 100 μl medium (IMDM). After a 5-hour coculture, the cells were harvested from the plates and transferred to V-bottom plates. Target cells that remained attached after the initial harvest were removed using trypsin and added to the V-bottom plate. The percentage of dead target cells and activation marker expression on γδ T cells were measured using flowcytometry.

### Flowcytometry

#### Antibodies used are listed in table 1

PBMCs or γδ T cells were washed with PBS prior to staining. Extracellular staining was performed by incubation with specific antibodies and LIVE/DEAD NEAR-IR (1:1000, Thermo Fisher Scientific) diluted in PBS for 30 min. on ice in the dark. The cells were washed twice with buffer (PBS/0.5% BSA) and either resuspended in PBS and directly analyzed or fixated using the BD Cytofix/Cytoperm fixation and permeabilization kit (BD Bioscience) according to the manufacturer’s protocol. For intracellular staining, cells were permeabilized and incubated with antibodies targeting IFN-γ, TNF-α or granzyme B diluted in permeabilization buffer for 30 min. on ice. Subsequently, the cells were washed twice and resuspend in PBS prior to analysis. Stained cells were analyzed or sorted on BD flow cytometers (LSR-II, Fortessa, FACSymphony or ARIA-II) and analyzed using FlowJo™ software version 10.9.0 (Ashland, OR: Becton, Dickinson and Company; 2023).

For analysis of γδ T cells in PBMCs, after isolation and stimulation, the cells were gated on single cells, time, NEAR-IR negative events and CD3, Vδ1 or Vδ2 positive events. For analysis of target cell killing, γδ T cell were distinguished using the gating strategy as described and remaining cells were gated for single cells and time before NEAR-IR evaluation representing dead cells.

### Statistical Analysis

Statistical testing was done by a student’s T-test or a one-way ANOVA followed by a Tukey’s multiple comparison test and are indicated in the figure legends. The statistical analysis was performed using GraphPad Prism version 10.0 for Mac OS (GraphPad Software, Boston, Massachusetts USA). Differences were considered significant when p≤0.05.

## Results

### Vδ2 T cells directly isolated from PBMCs can reach a 734-fold expansion on average during a 14-day culture period

Vδ2 T cells are the most prevalent γδ subset in the peripheral blood. To examine if Vδ2 T cells can be efficiently isolated directly from PBMCs and subsequently expanded, magnetic isolation was performed using mouse anti-Vδ2 TCR antibodies and anti-mouse IgG beads (MACS), followed by a FACS sort. MACS isolation generally yielded between 80-90% pure Vδ2^+^ T cells (of living cells) which could be improved to >99% with FACS sort (Fig. 1A, Suppl. Fig. 1A).

**Figure 1.**
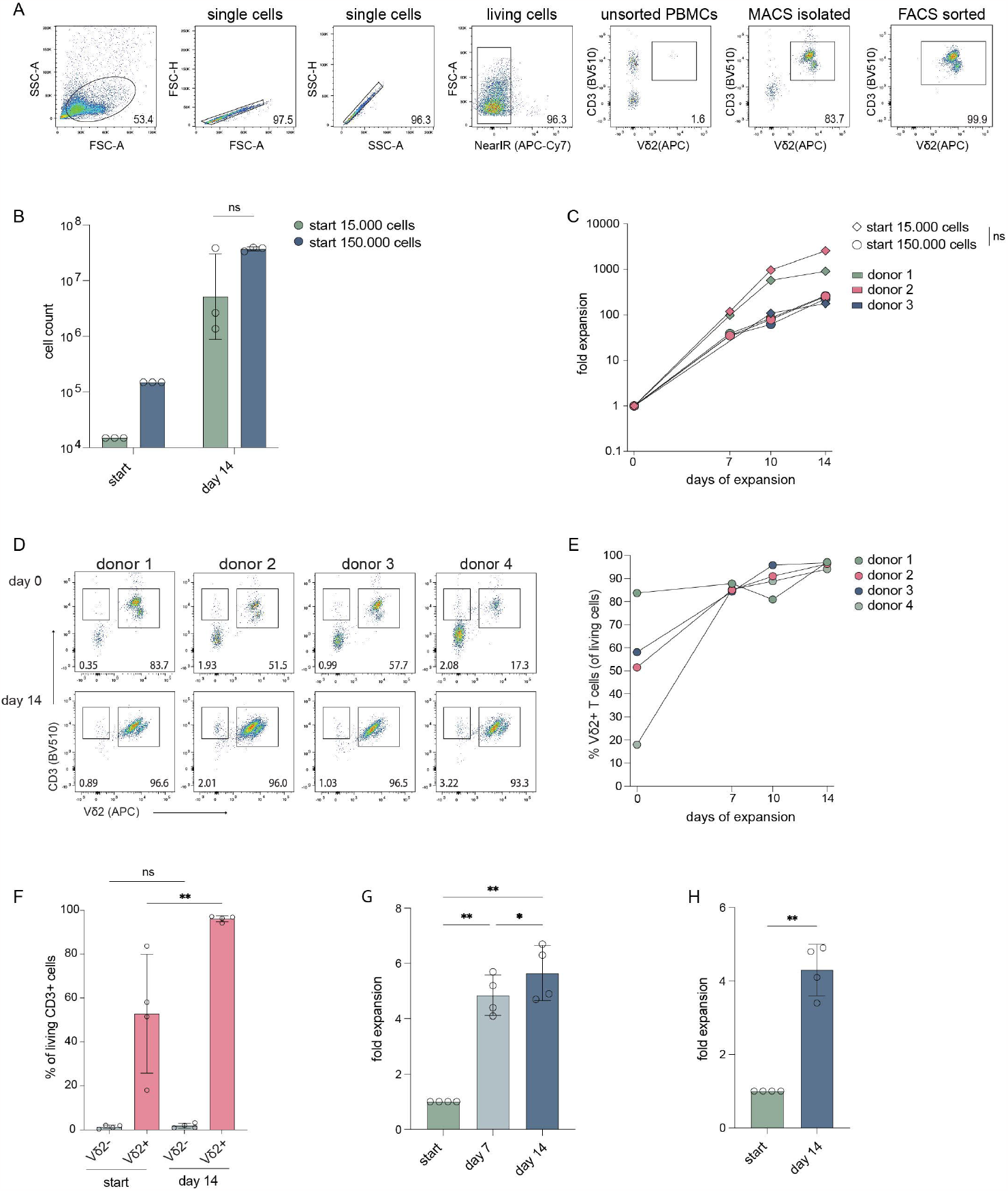
Cell count and expansion fold of Vδ2^+^ T cells after 14 days of expansion. The cells were isolated through mouse anti-human Vδ2 TCR and anti-mouse IgG bead MACS enrichment, with or without additional FACS sort, and expanded straight after isolation or previously expanded Vδ2^+^ T cells were submitted to a new round of expansion. (**A**) Representative flowcytometry plots and gating strategy showing the percentage of Vδ2^+^ T cells in unsorted PBMCs, after Vδ2 TCR specific MACS isolation and after an additional FACS sort for Vδ2^+^ T cells. (**B**) The number of Vδ2^+^ T cells used at the start of expansion, 15.000 and 150.000, and the number of cells yielded after the 14-day expansion (*n=3*). (**C**) The expansion fold of the Vδ2^+^ T cells for each donor of day 7, 10 and 14 relative to the start numbers, 15.000 and 150.0000, respectively (*n=3*). (**D**) Flowcytometry plots showing the percentage of the Vδ2^+^ cells after isolation (top row) using only the anti-Vδ2 TCR antibody combined with anti-mouse IgG beads and MACS separation, without FACS sort, and after a 14-day expansion period (bottom row) for four donors. (**E**) The percentage of Vδ2^+^ T cells after 7, 10 and 14 days of expansion and (**F**) the percentage of CD3^+^Vδ2^-^ T cells and CD3^+^Vδ2^+^ T cells before and after 14 days of expansion. (**G**) The expansion fold of Vδ2^+^ T cells that were expanded for 14 days that were, straight from culture, submitted to a new expansion culture of 14-days (*n=4*). (**H**) Expansion fold of Vδ2^+^ T cells that were previously expanded for 14 days after which they were cryopreserved for at least four weeks, thawed and submitted to a new expansion culture of 14 days (*n=4*). The data is shown as the mean and standard deviation of the donors and data from each donor represents the mean of triplicates. Data was analyzed by a one-way ANOVA followed by Tukey’s multiple comparisons test (**B, C, F, G**) or a student’s T-test (**H**). *P ≤0.05, **P ≤0.01, ns = not significant.

Culture of the cells was initiated at either 15.000 or 150.000 cells per well to determine the effect of cell seeding density on expansion. During expansion, the percentage of Vδ2^+^ T cells remained stable with a >98% purity with a minor reduction to approx. 95% at day seven likely because of TCR downregulation after activation (Fig. 1B. Suppl. Fig. 1A, B). Cultures which started with 15.000 cells reached cell numbers of 3.3-38.5 million cells at day 14 and cultures starting with 150.000 cells resulted in 33.9-39.5 million cells after 14 days (Fig. 1C). There was no significant difference in expansion between the different seeding densities (15.000 versus 150.000 cells) (Fig. 1C). Direct isolation targeting the Vδ2 TCR thus yields pure Vδ2^+^ T cells populations (>99%) which can successfully be expanded.

### FACS-sorted Vδ2 cells expand equally to non FACS-sorted cells

To assess if the process of FACS sorting following MACS enrichment affects the expansion potential of the isolated γδ T cells, the expansion capacities of isolated Vδ2^+^ T cells with or without FACS sort were compared from the same donors. The expansion of cells that were isolated using only MACS enrichment generated 12-43 million cells (Suppl. Fig. 2A, adjusted to percentage Vδ2^+^ cells, *n=4*) and displayed a similar expansion fold compared to cells from the same donor that were isolated through both MACS and FACS sort (Suppl. Fig. 2B, *n=3*). Thus, FACS sorting does not compromise the expansion capacity of Vδ2^+^ T cells.

**Figure 2.**
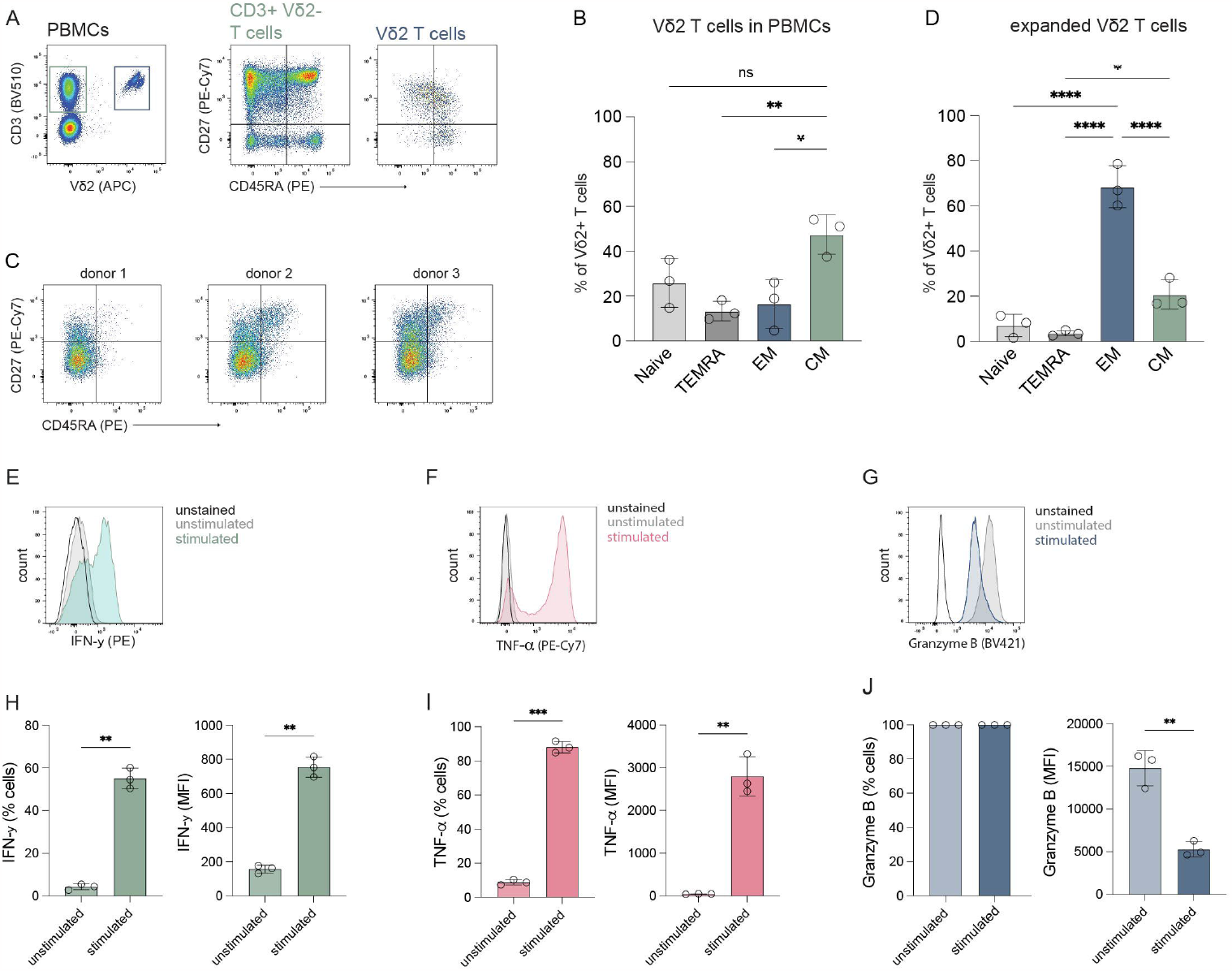
Differentiation phenotype and effector molecule production of expanded Vδ2^+^ T cells. Cells were stained for CD27 and CD45RA to determine their differentiation status and stimulated with PMA/Ionomycin to assess their potential to produce effector molecules IFN-γ, TNF-α and granzyme B. (**A**) Representative flowcytometry plots showing the frequency of CD3^+^Vδ2^-^ and CD3^+^Vδ2^+^ T cells in PMBCs (*left plot*) and the frequency of naïve (T_naive_, CD27^+^CD45RA^+^), terminally differentiated Effector Memory RA (T_EMRA_, CD27^-^CD45RA^+^), Effector Memory (T_EM_, CD27^-^CD45RA^-^) and Central Memory (T_CM_, CD27^+^CD45^-^) T cells for the CD3^+^Vδ2^-^ (*middle plot*) and CD3^+^Vδ2^+^ T cells (*right plot*) in PMBCs before isolation. (**B**) Summary of the percentage of T_naive_, T_EMRA_, T_EM_ and T_CM_ subsets in CD3^+^Vδ2^+^ T cells before isolation, as determined in **A**. (**C**) Flowcytometry plots of three donors showing the distribution of the T_naive_, T_EMRA_, T_EM_ and T_CM_ subsets in the total Vδ2^+^ T cell population after a 14-day expansion culture. Gating is based on freshly isolated CD3^+^ cells in total PMBCs from a reference donor. (**D**) Summary of the data in **C**. (**E-G**) Representative flowcytometry plots showing production of IFN-γ, TNF-α of granzyme B by Vδ2^+^ T cells stimulated with PMA/Ionomycin for 1.5 hour or unstimulated. (**H-J**) Summary of the production of IFN-γ, TNF-α of granzyme B by Vδ2^+^ T cells stimulated with PMA/Ionomycin for 1.5 hour or unstimulated. The percentage and MFI of the cytokine and granzyme B positive cells is depicted. Data are shown as the mean and standard deviation of three donors and the data of each donor represents the mean of triplicates. Data were analyzed by a one-way ANOVA followed by Tukey’s multiple comparisons test (**B, D**) or a student’s T-test (**H-J**). *P ≤0.05, **P ≤0.01, ***P ≤0.001, ****P ≤0.0001. ns = not significant.

### The expansion of Vδ2 T cells is not compromised by outgrowth by other cell types

Given the generic character of the feeder-based expansion protocol, other T cell subsets may potentially overgrow the γδ T cells during the expansion phase. To investigate whether the purity of the Vδ2^+^ T cells could be compromised by the contamination of αβ T cells or other cell types, less pure Vδ2^+^ T cells isolated using direct targeting MACS isolation (*n=4*) or pan γδ T cells isolated using untouched MACS isolation (*n=2*) without FACS sort were expanded and significant co-expansion of other cells was monitored.

After direct Vδ2^+^ targeting MACS isolation, the percentage of CD3^+^Vδ2^+^ T cells ranged between 17-84% (μ=53%) (Fig. 1D-F) and between 0.4-2% (μ=1%) of the isolated cells was CD3^+^Vδ2^-^. Most Vδ2 negative cells that remain after MACS isolation were not T cells (CD3^-^). The percentage of CD3^+^Vδ2^+^ T cells significantly increased to 93-97% (μ=96%) after a 14-day expansion period (Fig. 1E). The CD3^+^Vδ2^-^ cell population expanded alongside the CD3^+^Vδ2^+^ T cells during the 14-day culture though their relative presence did not significantly increase (Fig. 1D, F). The CD3^-^ cells were no longer present after the expansion (Fig. 1D).

From two donors, γδ T cells were isolated from PBMCs using the TCRγ/δ+ T Cell Isolation Kit (Miltenyi), which comprised of 49% and 80% Vδ2^+^ T cells. Expansion of these γδ T cells yielded 20-27 million cells of which 76-96% were Vδ2^+^ T cells and 3-20% are CD3^+^Vδ2^-^ cells (Suppl. Fig. 2C-E). Altogether, this indicates that - during the feeder-based expansion protocol - the purity of Vδ2^+^ T cells is not compromised by overgrowth by other cells, such as αβ T cells.

### Previously expanded Vδ2 T cells can be expanded further using this PHA based TCR triggering culture method

To examine if Vδ2^+^ T cells can be expanded further after initial expansion, Vδ2^+^ T cells were submitted to a new 14-day expansion culture straight from the first 14-day expansion. The Vδ2^+^ T cells (*n=4*) reached a 4.1-5.7-fold expansion (μ=5) after the first seven days and a 4.7 to 6.7-fold expansion (μ=6) after the second full 14-day expansion period (Fig. 1G).

To investigate whether 14-day expanded Vδ2^+^ T cells can be expanded further after cryopreservation, frozen 14-day expanded Vδ2^+^ T cells were thawed and submitted to a new 14-day expansion. Frozen, previously expanded Vδ2^+^ T cells (*n=4*) showed a 4.3 to 4.9-fold expansion after 14 days (Fig. 1H).

Although reduced compared to the first expansion, previously expanded Vδ2^+^ T cells can be expanded further using the expansion protocol described in this report.

### The Vδ2 T cells show an effector phenotype after expansion

To assess the phenotype and functionality of the expanded Vδ2^+^ T cells, the cells were immunophenotyped before and after expansion using surface expression of CD27 and CD45RA and evaluated for their ability to produce effector molecules.

Before expansion, 38-54% (μ=48%) of the Vδ2^+^ T cells were primarily of a central memory (T_CM_; CD27^+^CD45RA^-^) phenotype followed by a naïve phenotype (T_naive_; CD27^+^CD45RA^+^), with 15-36% (μ=26%) an effector memory phenotype (T_EM_; CD27^-^CD45RA^-^) with 5-26% (μ=17%) (Fig. 2A, B). The percentage of terminally differentiated effector memory RA (T_EMRA_; CD27^-^CD45RA^+^) cells was 9-18% (μ=13%). After expansion, most Vδ2^+^ T cells exhibited a T_EM_ phenotype which comprised 60-79% (μ=69%) of the cells, followed by a T_CM_ phenotype containing 17-28% (μ=21%) of the cells (fig. 2C, D). The T_naive_ and the T_EMRA_ subsets contained the fewest cells with 2-11% (μ=7%) and 3-5% (μ=4%), respectively.

To further asses effector potential, expanded Vδ2^+^ T cells were stimulated with PMA/Ionomycin for 1.5 hours. Up to 60% of γδ T cells produced IFN-γ and 84-92% produced TNF-α (Fig. 2E, F, H, I). Production of granzyme B was present in most cells before stimulation (>95%) and cells remained positive upon activation, but showed a significant decrease in MFI for granzyme B staining, possibly indicating early degranulation (Fig. 2G, J).

These data show that positively isolated Vδ2^+^ T cells expanded with feeder cells, PHA, IL-2, IL-7 and IL-15 predominately have an effector/central memory phenotype with the capacity to produce IFN-γ, TNF-α and granzyme B.

### The Vδ2 T cells expanded with feeder cells, PHA, IL-2, IL-7 and IL-15 effectively kill target cells

The capacity to induce anti-tumor cytotoxicity by positively isolated and expanded Vδ2^+^ T cells was assessed in co-cultures with cell lines derived from different tumor types; WiDr cell line (colon adenocarcinoma), WM9 cell line (melanoma) and the HAP1 cell line (chronic myelogenous leukemia) which, due to its near-haploidy, is a highly useful cell line for genetic manipulation/gene knock-out approaches in future studies on mechanistic pathways (52). Vδ2^+^ T cells from three different donors effectively killed 35-40% WiDr cells (μ=39%), 15-20% WM9 cells (μ=17%) and 25-34% HAP1 cells (μ=30%) (Fig. 3A-D). Pretreatment with PAM significantly improved HAP1 cell killing by Vδ2^+^ T cells but not of WiDr and WM9 cells (Fig. 3D). The activation state of expanded Vδ2^+^ T cells via CD25, CD69 and CD137 was also assessed in cocultures with WiDr, WM9 and HAP1 tumor cells (Fig. 3E-H). Surface expression of CD25 and CD69 was significantly increased on Vδ2^+^ T cells cultured with WiDr cells, indicating that these target cells most strongly activated the Vδ2^+^ T cells (Fig. 3E-G).

**Figure 3.**
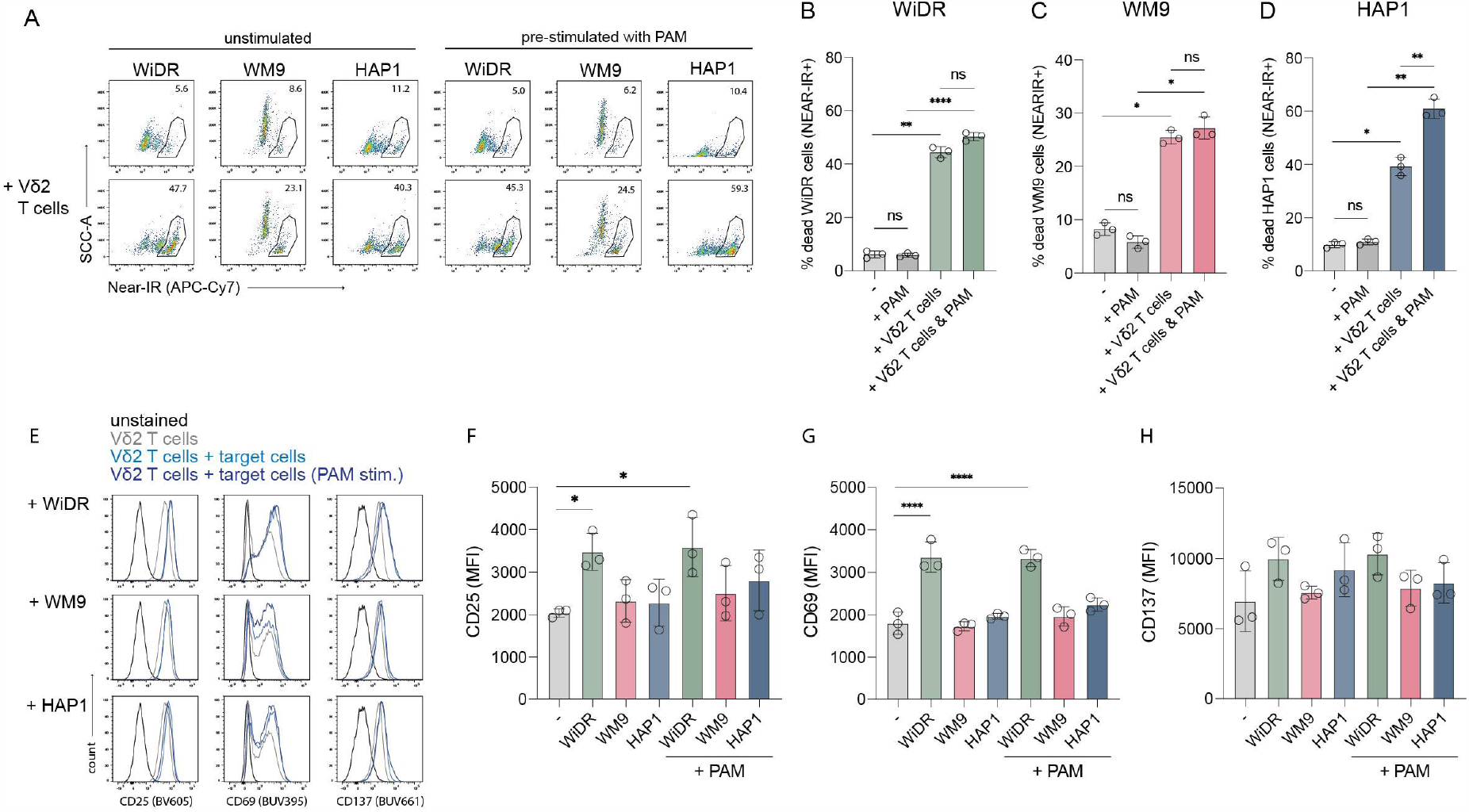
Coculture of freshly expanded Vδ2^+^ T cells with target cell lines WiDr, WM9 and HAP1. 15.000 cells were plated one day prior to the coculture with or without PAM prestimulation. An effector:target ratio of 5:1 was used in a 5-hour coculture. (**A**) Representative flowcytometry plots showing the frequency of dead (Near-IR+) target cells only (*top row*) and target cells cocultured with Vδ2^+^ T cells (*bottom row*), with or without pretreatment with PAM. (**B-D**) Summary of the percentage of dead target cells, WiDr, WM9 and HAP1, only (-) or with PAM treatment and/or addition of Vδ2^+^ T cells. (**E**) Representative histograms plots showing the cell surface expression of CD25, CD69 and CD137 of Vδ2^+^ T cells only or cultured with indicated target cell lines either prestimulated with PAM or not. (**F-G**) Overview of the MFI of CD25, CD69 and CD137 of Vδ2^+^ T cells only (-) or cultured with indicated target cell lines either prestimulated with PAM or not. Gates were set using unstained Vδ2^+^ T cells. Data are shown as the mean and standard deviation of the donors (*n=3*). Data of each donor represents the mean of triplicates. Data were analyzed by a one-way ANOVA followed by Tukey’s multiple comparisons test. *P ≤0.05, **P ≤0.01, ****P ≤0.0001. ns = not significant.

Altogether, these data showed that the Vδ2^+^ T cells expanded with feeder cells, PHA, IL-2, IL-7 and IL-15 could effectively kill tumor cells.

### Direct targeted MACS isolation combined with FACS sort and this expansion protocol can be used to expand both Vδ1 and Vδ2 subsets

To investigate if the direct isolation protocol can be used to purify and isolate Vδ1^+^ T cells and if their expansion would be equal to that of Vδ2^+^ T cells, both Vδ1^+^ and Vδ2^+^ T cells were isolated from the same PBMC donors and to compare their expansion competence.

Direct positive isolation of Vδ1^+^ and Vδ2^+^ T cells by magnetic beads (MACS) yielded approx. 90% Vδ1^+^ or Vδ2^+^ pure populations, with purity being increased to > 99% purity by additional FACS sort (Fig. 4A). Over the culture period of 14 days, the two subsets expanded within a similar range through the first week (day 7) with an expansion of 29-fold for Vδ1^+^ and 19-fold for Vδ2^+^ T cells on average (Fig. 4B). At day 14, expansion of both subsets was high, with expansion of the Vδ2^+^ subset exceeding of the Vδ1^+^ subset with an average of 1248-fold over a 380-fold for Vδ1^+^ T cells (Fig. 4B).

**Figure 4.**
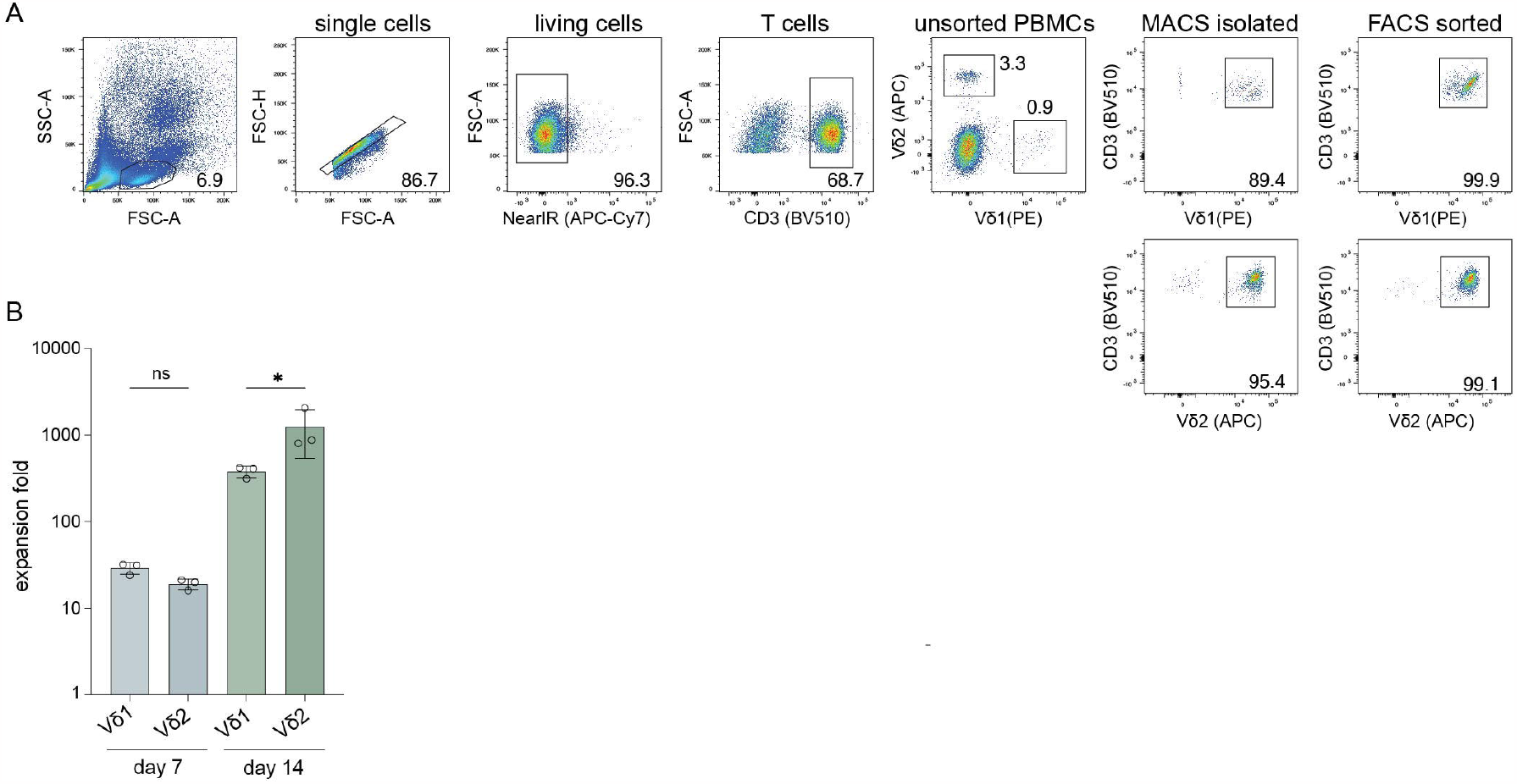
Comparison of the expansion competence of Vδ1^+^ T and Vδ2^+^ T cells of the same donors starting at 150.000 cells per well. The cells were isolated through mouse anti-human Vδ1 TCR or Vδ2 TCR and anti-mouse IgG bead MACS enrichment and further purified using FACS sort. (**A**) Representative flowcytometry plots of the percentage of Vδ1^+^ T cells and Vδ2^+^ T cells in PBMCs, after Vδ1^+^ or Vδ2^+^ T cells specific MACS isolation and after an additional SORT of Vδ1^+^ or Vδ2^+^ T cells before expansion. (**B**) The expansion ratio of Vδ1^+^ compared to Vδ2^+^ T cells after seven days and at the end of the expansion culture on day 14. Ratios were calculated using the cell count on either day 7 or day 14 and compared to the start number (150.000). Data were analyzed using a one-way ANOVA followed by Tukey’s multiple comparisons test (**A**). *P ≤0.05. ns = not significant.

This isolation and expansion method can thus be applied to both Vδ1^+^ and Vδ2^+^ T cells to yield high cell numbers for downstream use.

### Expansion of Vδ1 and Vδ2 T cells can be achieved from low starting numbers

γδ T cells are of great interest as a means of cellular therapy to treat cancer and understanding the functionality of these cells within tumors is important for the development and optimization of future therapies. However, the absolute amount of γδ T cells isolated from TILs in primary tumors is often very low which complicates further research. Therefore, the applicability of this expansion protocol was evaluated to expand Vδ1^+^ and Vδ2^+^ T cells from very low starting numbers.

Expansion of directly isolated Vδ1^+^ and Vδ2^+^ T cells (purity >95%) from three donors with an initial seeding density of 150 cells per well showed expansion ratios of >1000 fold at day 17 of expansion (Fig. 5). After a 24-day culture period, the cells had reached a plateau in expansion. There was no significant difference observed between the Vδ1^+^ and Vδ2^+^ T cells.

**Figure 5.**
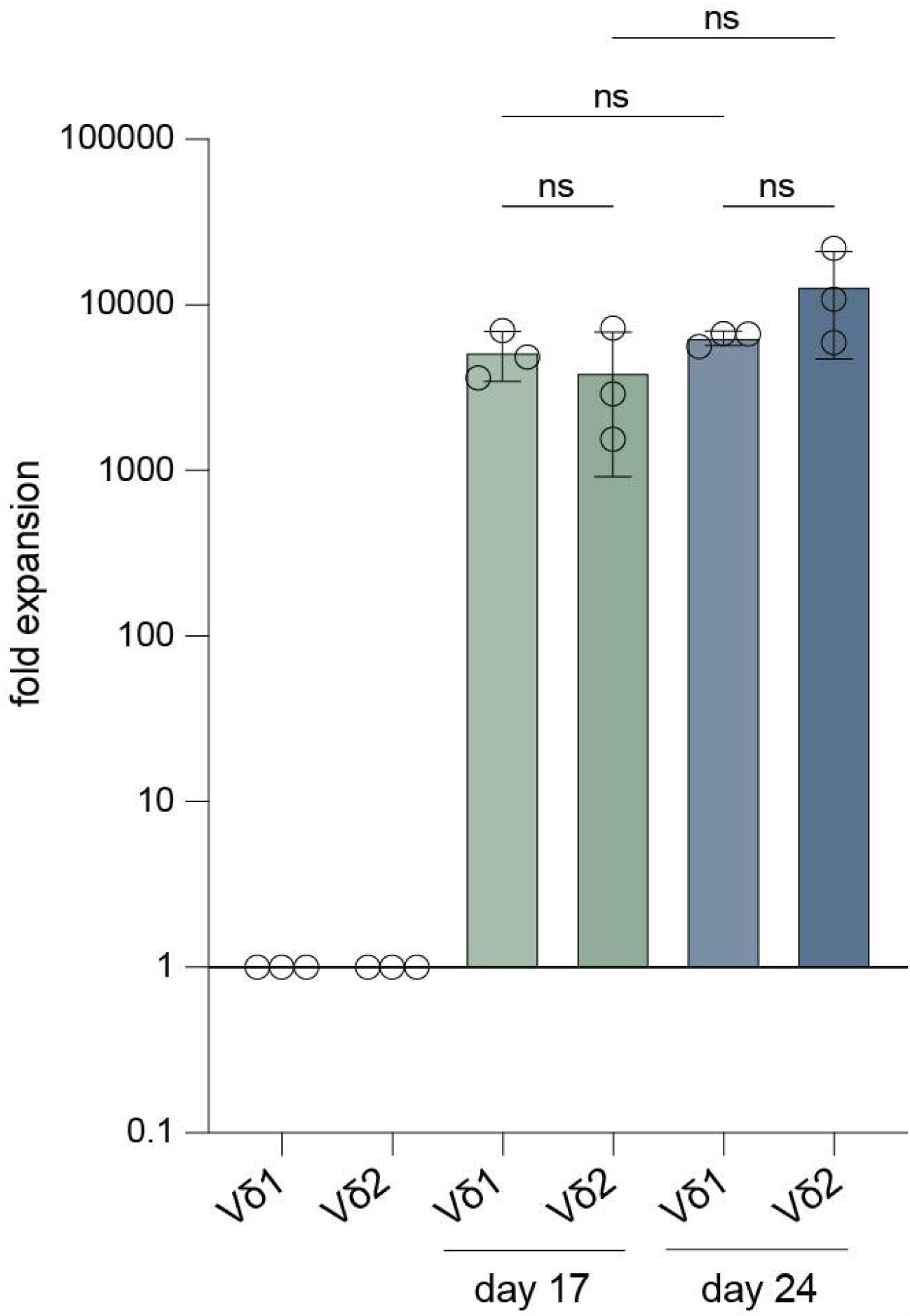
Expansion fold of Vδ1^+^ and Vδ2^+^ T cells starting from low cell numbers. Vδ1^+^ and Vδ2^+^ T cells were isolated from three donors using mouse anti-human Vδ1 or Vδ2 TCR and anti-mouse IgG bead MACS enrichment, followed by FACS sort and 150 cells/well were expanded for 24 days. The expansion fold of Vδ1^+^ and Vδ2^+^ T cells is shown after 17 days and at the end of the expansion culture on day 24 (*n=3*). Ratios were calculated using the cell count on either day 17 or day 24 and compared to the start number (150). Data were analyzed using a one-way ANOVA followed by Tukey’s multiple comparisons test. Ns = not significant.

Thus, this isolation and expansion method is also effective with low amounts of starting material and might potentially be successful in expanding tumor tissue-derived γδ T cells.

## Discussion

We described a methodology to directly isolate γδ T cells from PBMCs using MACS enrichment followed by FACS-based sorting, generating >99% pure γδ T cells. These cells effectively expand and produce effector molecules IFN-γ, TNF-α and granzyme B upon stimulation. In addition, the expanded γδ T cells successfully kill cell lines of different tumor types. More importantly, we show that this method can be applied to expand γδ T cells from low cell numbers (i.e., 150 cells) and may therefore provide a tool to study γδ T cells *ex vivo*.

The here described expansion protocol is based on directly isolating γδ T cells from freshly isolated PBMCs in contrast to most reported expansion protocols which generally deplete αβ T cells and/ or CD56 positive cells (*50, 51, 53*–*56*). A disadvantage of those methods is that also a fraction of γδ T cells can be removed during this depletion step since they can express CD4, CD8 and CD56 (*9, 57*–*61*). On the other hand, the efficiency of depleting other immune cell subsets is often not reported and, in contrast to FACS sort, this depletion is not fully sufficient to remove non-γδ T cells (*50*). As a consequence, αβ T cells, NK cells and others such as B cells and monocytes might contaminate the γδ T cell cultures (*51*), which is not preferable because their contribution can result in misrepresentation of the functionality of γδ T cell in *in vitro* assays.

It is not completely clear whether having αβ T cells present during the expansion period itself is a benefit or a harm for the expansion of γδ T cells since these cells thrive on the cytokines included in culture. Though, the data on cells isolated through MACS bead separation without FACS sort presented here showed that a small fraction of non-Vδ2 T cells do not overtake γδ T cells during expansion which indicates that the presence of other T cells does not result in an adverse effect on γδ T cell expansion. In addition, the data shown in this report supports that other cell types such as NK cells, B cells and monocytes most likely do not grow out using this MACS isolation and expansion method.

During cultures including other cell types, the γδ T cells may receive survival signals from these cell types that facilitate the expansion. One study suggests that δγ T cells expand better when the other T cells are in the same culture (*62*). The culture method proposed in this report includes feeder cells comprised of irradiated PBMCs and EBV transformed B cells which may account for these signals. Evaluation of this culture method with addition or substitution of specific (feeder) cells in future may provide more insights into benefit of the other cell types during expansion.

This study is not the first to include a FACS sort for γδ T cell isolation. The added benefit of the protocol described here lies in the combination of MACS isolation and FACS sort, which makes it a very efficient isolation method in terms of yield and consumption of time. Cho and colleagues uses positive selection of Vδ2 T cells solely through FACS sort prior to expansion using K562 feeder cells, anti-CD3/28 and IL-2 (*63*). Merely a FACS sort is likely to obtain a pure γδ T cell population from PBMCs, however, FACS sorting γδ T cells (0.5-5% of CD3^+^ T cells) is very time-consuming. The pre-enrichment of γδ T cells by MACS isolation (purity γδ T cells of total cells: 80-90%) as described here, greatly reduces the time necessary to FACS sort the cells.

The expansion ratio for Vδ1 and Vδ2 T cells using the protocol described here (starting at 150.000 cells) range between 200 to 2000-fold. These cells can be expanded for a second time, either from culture or cryopreservation, allowing the generation of more cells and repeat experiments without the need of new γδ T cell isolation.

Ferry and colleagues depleted αβ T cells and CD56^+^ NK cells from PBMCs and stimulated γδ T cell expansion using OKT-3 (anti-CD3) in combination with IL-15 after which they depleted Vδ2 T cells and focused on generating multi-applicable Vδ1 T cells (*50*). Interestingly, when they compared expansion of the Vδ1 T cells with OKT-3 + IL-15 to PHA + IL-2 or IL-7 they observed a better yield with OKT-3 compared to PHA conditions. Moreover, the OKT-3 activated Vδ1 T cells showed higher expression of activation markers CD69 and NKG2D. However, expansion factors reached with these protocols ranged between 10 to 48-fold using OKT-3 + IL-15 within 20-30 days, while we reached a >300-fold expansion for Vδ1 T cells with PHA and feeder cells in 14 days. This difference could be caused by that the authors may have used a different PHA while the PHA (HA-16) used in this report has repeatedly been confirmed to have high efficiency (*52, 64, 65*).

We were able to expanded 150 Vδ1 and Vδ2 T cells to 0.8-3.3 million cells in 3.5 weeks, corresponding to an expansion ratio between 5.600-22.000. De Vries and colleagues FACS sorted between 168 and 3775 γδ T cells derived from colon cancer tissue which were expanded for 3-4 weeks using PHA, IL-2 and IL-15, reaching a 2000 to 170.000-fold expansion (*66*). An explanation for the higher expansion ratios achieved in that study may be a difference in activation state of the isolated cells γδ T cells from tumor tissue. Another explanation is the IL-2 concentration of 1000 units/ml used while the expansion medium used here contained 120 units/ml IL-2. Whether the protocol described here can be further optimized in terms of cytokine concentrations or inclusion of other immune cell activating cytokines such as IL-4, IL-18 and IL-21, used in several other expansion protocols, needs further investigation.

Functionality of expanded γδ T cells was confirmed by lysis of WiDr, WM9 and HAP1 tumor cells. Interestingly, and in line with our findings, Correira and colleagues showed cytotoxicity towards leukemia-derived cell lines by PHA-expanded γδ T cells. These PHA-expanded γδ T cells were also superior over bromohydrin pyrophosphate (HMB-PP)-expanded γδ T cells in anti-tumor cytotoxicity (*67*). This combined with our data suggests that TCR triggering via crosslinking by PHA generates γδ T cells with a higher killing capacity compared to phosphoantigen stimulated expansion.

Most reported expansion methods only focus on Vδ2 T cells while Vδ1 T cells also have pro- and anti-tumor potential which renders these cells highly relevant to investigate (*38, 68*). For example, it has been found that the Vδ1 T cells subset is most prevalent in TILs extracted from melanoma (*30, 69*). Importantly, these cells were reactive towards both allogeneic and autologous melanoma (*69*). In addition, Vδ1 T cells showed robust cytotoxicity against colorectal cancer (CRC), hepatocellular cancer (HCC) and leukemia *in vitro* (*67, 70, 71*). In contrary, it has been shown that IL-17 produced by Vδ1 cells recruits myeloid derived suppressor cells and that they suppress αβ T cell function and DC maturation (*72, 73*). Therefore, it is important to expand both subsets as is possible with the direct MACS and FACS sort isolation described in this study.

The Vδ1 or Vδ2 subsets are most studied in γδ T cells research while Vδ3 T cells are also found in the blood and in tumors (*66*). Whether this subtype can also be expanded to generate large quantities of cytotoxic γδ T cells using the protocol described here has to be explored.

## Supporting information

Protocol

## Supplementary figures

**Figure S1**. The purity of Vδ2^+^ T cells isolated with mouse anti-human Vδ2 TCR and anti-mouse IgG bead MACS enrichment followed by a Vδ2^+^ FACS sort during the 14-day expansion. **(A)** Flowcytometry plots of three donors showing the percentage of Vδ2^+^ T cells on day 0 (start), 7, 10 and 14. (**B**) Summary of the percentage of the data in **A** (*n=3*). Data of the 150.000 start condition is shown. Data was analyzed using a one-way ANOVA followed by Tukey’s multiple comparisons test. *P ≤0.05, ns= not significant.

**Figure S2**. Expansion of CD3^+^Vδ2^+^ and CD3^+^Vδ2^-^ cells isolated using mouse anti-human Vδ2 TCR and anti-mouse IgG bead MACS enrichment, with or without further purification using a FACS sort, or using untouched MACS isolation with the TCRγ/δ+ T Cell Isolation kit. (**A**) Cell count of Vδ2^+^ cells isolated using anti-Vδ2 MACS bead separation only after 7, 10 and 14 days of culture to expand the cells (*n=4*). Counts are adjusted to the purity of the cells on the indicated days. (**B**) Comparison of expansion fold of Vδ2^+^ cells isolated using anti-Vδ2 MACS bead separation only or with an additional FACS purification (*n=3*). The cells from the same donors are compared. Counts that are used to calculate the expansion ratio are adjusted to the purity of the cells on the indicated days. (**C**) Flowcytometry plots illustrating the percentage of CD3^+^Vδ2^-^ and CD3^+^Vδ2^+^ of cells isolated with the TCRγ/δ+ T Cell Isolation Kit after 14 days expansion (*n=2*). (**D**) Cell count of CD3^+^Vδ2^+^ T cells isolated using the TCRγ/δ+ T Cell Isolation Kit before and after 14-days expansion (*n=2*) (**E**) Percentage of CD3^+^Vδ2^+^ T cells in the total population isolated using the TCRγ/δ+ T Cell Isolation Kit before and after 14-days expansion (*n=2*). Data were analyzed by using a student’s T-test. *P ≤0.05, ns= not significant.

## Notes

### Competing Interest Statement

The authors have declared no competing interest.

