## Supplementary material for "Isolation and expansion of pure and functional γδ T cells": Protocol

### Step by step protocol

#### Medium

*γδ T cells basic medium*
Basic γδ T cell medium is composed of IMDM supplemented with 5% FCS, 5% human serum and 1% pen/strep.

*γδ T cells expansion start medium*
For the initial expansion, the γδ T cells basic medium is completed with 1.0 μg/ml PHA, 20 ng/ml IL-7, 20 ng/ml IL-15, 120 units/ml IL-2 and feeder cells: 100.000 irradiated EBVLCLs (50 Gy) of at least two different sources and 1 million irradiated PBMCs (30 Gy) of at least three different donors per ml of γδ T cell medium. In total, 150.000 γδ T cells are resuspended per ml of complete expansion medium and plated in a 24-well plate at 1 ml per well (table 2).

*Table 2.* Overview of the components of γδ T cells expansion start medium

| *reagent* | *Final concentration per ml basic medium* |
|---|---|
| IL-2 | 120 U |
| IL-7 | 20 ng |
| IL-15 | 20 ng |
| PHA | 1.0 μg |
| feeder cells | 100.000 EBVLCLs (2 sources), irradiated with 50 Gy<br>1 million PBMCs (3 donors), irradiated with 30 Gy |

*γδ T cell expansion complete medium*
For the addition of medium at day 3 and day 10, make a 5x concentrated premix of basic γδ T cell medium supplemented with cytokines to reach a final concentration of 100 ng/ml IL-7 (final 20 ng/ml), 100 ng/ml IL-15 (final 20 ng/ml) and 600 units/ml IL-2 (final 120 units/ml).

#### Isolation of γδ T cells from PBMCs (I)

A buffy coat generates approx. >1x$10^9$ PBMCs containing 0.5-5% Vδ2 T cells. This protocol focuses on isolating approx. 300.000-1.000.000 γδ T cells from 350 million PBMCs. This isolation method uses mouse anti-γδ TCR and anti-mouse IgG beads followed by MACS isolation and a FACS sort which results in the highest purity. Samples are taken after different isolation steps to assess purity of the cells (part II).

Preparation: Cool down the centrifuge to 4°C, prewarm the γδ T cell basic medium and prepare cold PBS/0.02% human albumin/2mM EDTA (pH8).

I. For the isolation of γδ T cells (Vδ1 and/or Vδ2), collect at least 350 million PBMCs in a 50 ml Falcon tube and perform a spin of 5 min. at 300G and 4°C. Take two samples of 20-50 μl from the remaining PBMCs, transfer these cells into a V-bottom plate and store at 4°C for the purity validation in ***part II.***
*note: when a FACS sort is not performed, collect separate PBMC samples for each subset.*

II. Prepare antibody solutions of 25 μl conjugated mouse-anti-human Vδ1 and/or Vδ2 in 500 μl cold PBS. It is recommended to use a maximal dilution of 1:50 of the conjugated anti Vδ1 and/or Vδ2 antibody. Add the antibody solution to the designated PBMCs, resuspend gently and incubate for 30 min. at 4°C. Resuspend gently after halfway the incubation period.

III. *note: when multiple γδ subsets are isolated from the same PBMC population, use different conjugated fluorochromes to distinguish the subsets with FACS sort.*

IV. Wash the cells by adding cold PBS/0.5% BSA up to a volume of 45 ml. Spin the cells for 5 min. at 300G and 4°C. Resuspend the cells in 1 ml sterile PBS and top up to 35 ml.

V. Take a small sample of 20 µl (approx. 200.000 PBMCs), transfer these cells to the V-bottom plate (step 1) and store at 4°C for the purity validation in ***part II***.

VI. Spin the Falcon tube with the stained cells 10 min. at 300G and 4°C.

VII. Resuspend the cells in 2.8 ml cold PBS/0.5% BSA. Add 700 µl anti-mouse IgG beads, resuspend gently and incubate for 15 min. at 4°C. Vortex gently halfway the incubation period.

VIII. Wash the cells by adding cold PBS/0.5% BSA up to a volume of 45 ml. Spin the cells for 10 min at 300G and 4°C. Resuspend the cells in 1 ml sterile PBS and repeat this step.

IX. Resuspend the cells in 1-2 ml cold PBS/0.5% BSA.

X. Perform the magnetic separation (MACS) according to the protocol provided by Miltenyi. Using an LS column is recommended to obtain optimal purity.

XI. Collect the flowthrough in a 15 ml Falcon tube, and perform a spin of 5 min. at 300G and 4°C.

XII. Resuspend the cells in 1 ml of basic γδ T cell medium and count the cells.

XIII. Take a small sample of a few thousand cells, transfer these cells to the V-bottom plate and store at 4°C for ***part II***.

XIV. Continue with part II for validation of viability and purity or part III for culturing of the cells.

#### γδ T cell viability and purity check (II)

For optimal expansion of γδ T cells and to minimize interference of other immune cell subsets in subsequent functional essays, aim for a purity of >85%.

*Table 3.* collected samples for the viability and purity check.

| sample | condition | source |
|---|---|---|
| PBMCs | unstained | step I.I |
| PBMCs | Stained for CD3, anti-human Vδ1 and/or Vδ2 and NEARIR | step I.V |
| Isolated γδ T cell | Stained for CD3, anti-human Vδ1 and/or Vδ2 and NEARIR | step I.XIII |

I. Add 150 µl cold PBS to the wells containing the cells and perform a spin of 2 min at 500G and 4°C.

II. Meanwhile, prepare a staining solution containing anti-human CD3 and NEARIR in PBS.

III. Resuspend the cells – except for the unstained PBMC control - in 35-50 µl staining solution and incubate for 20-30 min at 4°C.

IV. Add 150 µl cold PBS and perform a spin of 2 min at 500g and 4°C. Repeat this step.

V. Resuspend the cells in PBS and analyze the viability and purity of the γδ T cells using flowcytometry.

VI. To analyze the viability and purity of the isolated cells, the gating strategy as shown in Fig. 1 can be used.

When the cells have a purity <85% or a higher purity is preferred, an FACS sort can be performed (figure 1). The γδ T cells are stained for the Vδ1 and/or Vδ2 receptor and can therefore be sorted straight after the MACS isolation and do not require re-staining. Cells obtained through an additional FACS sort usually are >98% pure and have equal capacity to expand (figure 5).

#### γδ T cell culture and expansion (III)

*Day 0*

I. Take the γδ T cells isolated in *part I* and perform a spin of 5 min at 300G at RT and resuspend the cells in prewarmed complete γδ T cell expansion medium (*see materials*) at a concentration of 150.000 γδ T cells per ml.

II. Distribute the cells in a 24-well plate at 1 ml per well.
III. Culture the cells in the incubator at 37°C and 5% $CO_2$.

*Day 3*

Add 250μl of the 5x concentrated 'γδ T cells expansion complete medium' per well.
*Note: In case of another volume, adjust the cytokine concentration to obtain a final concentration of 120 units/ml IL-2, 20 ng/ml IL-7 and 20 ng/ml IL-15.*

*Day 7*

I. Harvest and pool the cells from the 24-well plate and count the cells.
II. If desired, a small sample can be taken to analyze the viability and purity using flowcytometry.
III. Add γδ T cell basic medium to obtain a concentration of 0.5 x$10^6$ cells per ml.
IV. Add cytokines: 120 units/ml IL-2, 20 ng/ml IL-7 and 20 ng/ml IL-15.
V. Plate 1 ml per well in a 24-well plate.
VI. Culture the cells in the incubator at 37°C and 5% $CO_2$.

*Day 10*

I. Harvest and pool the cells from the 24-well plate and count the cells.
II. Add γδ T cell basic medium to obtain a concentration of 0.5 x$10^6$ cells per ml.
III. Add cytokines: 120 units/ml IL-2, 20 ng/ml IL-7 and 20 ng/ml IL-15.
IV. Plate 1 ml per well in a 24-well plate.
V. Culture the cells in the incubator at 37°C and 5% $CO_2$.

*Day 14*

Harvest and pool the cells from the 24-well plate and count the cells. The cells can be used directly for (functional) analysis or can be frozen. It is recommended to take a small sample and analyze the viability and purity of the expanded cells before use.
