## Supplementary figures and images for "Isolation and expansion of pure and functional γδ T cells"

### supplemental figure 1

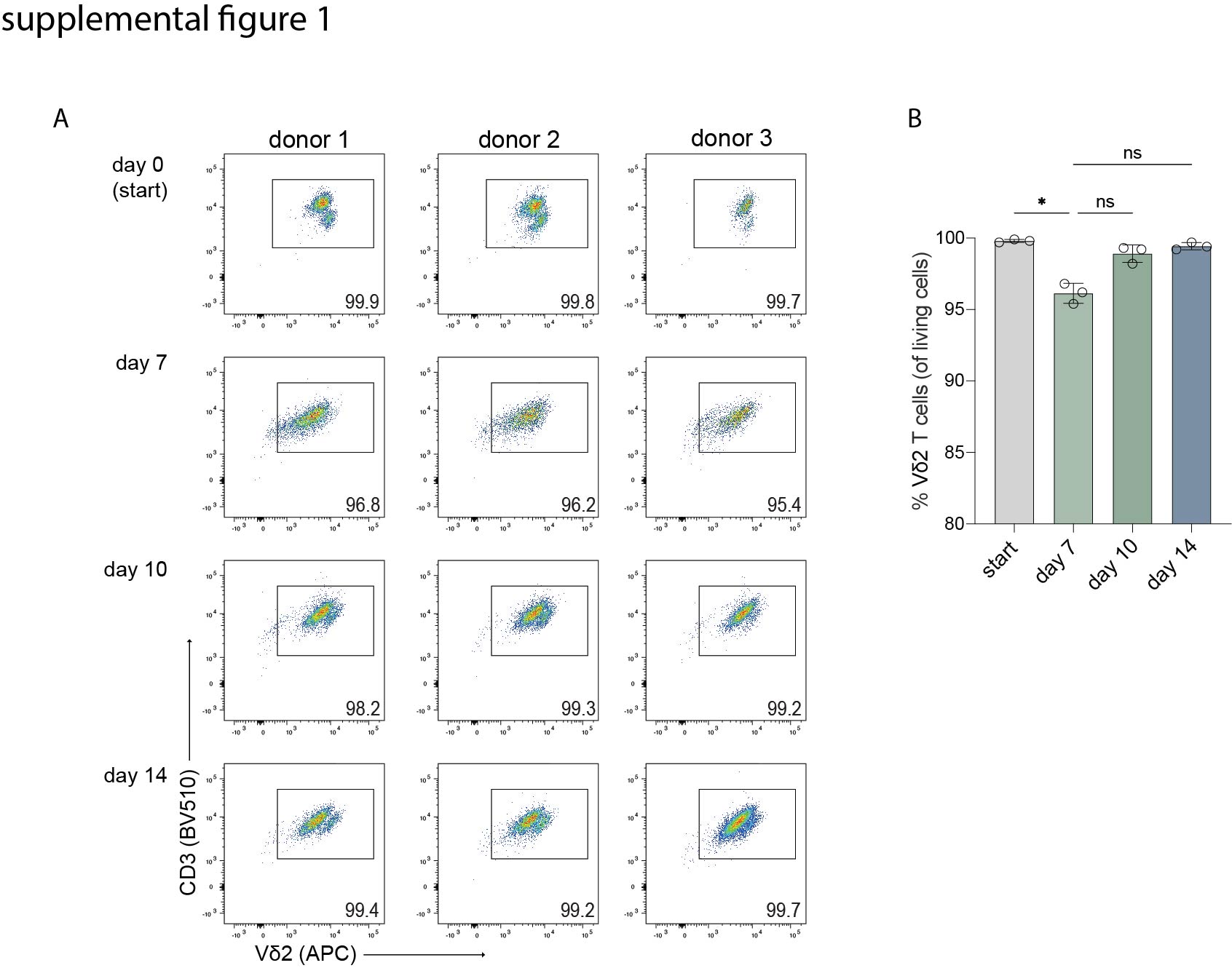

### supplemental figure 2

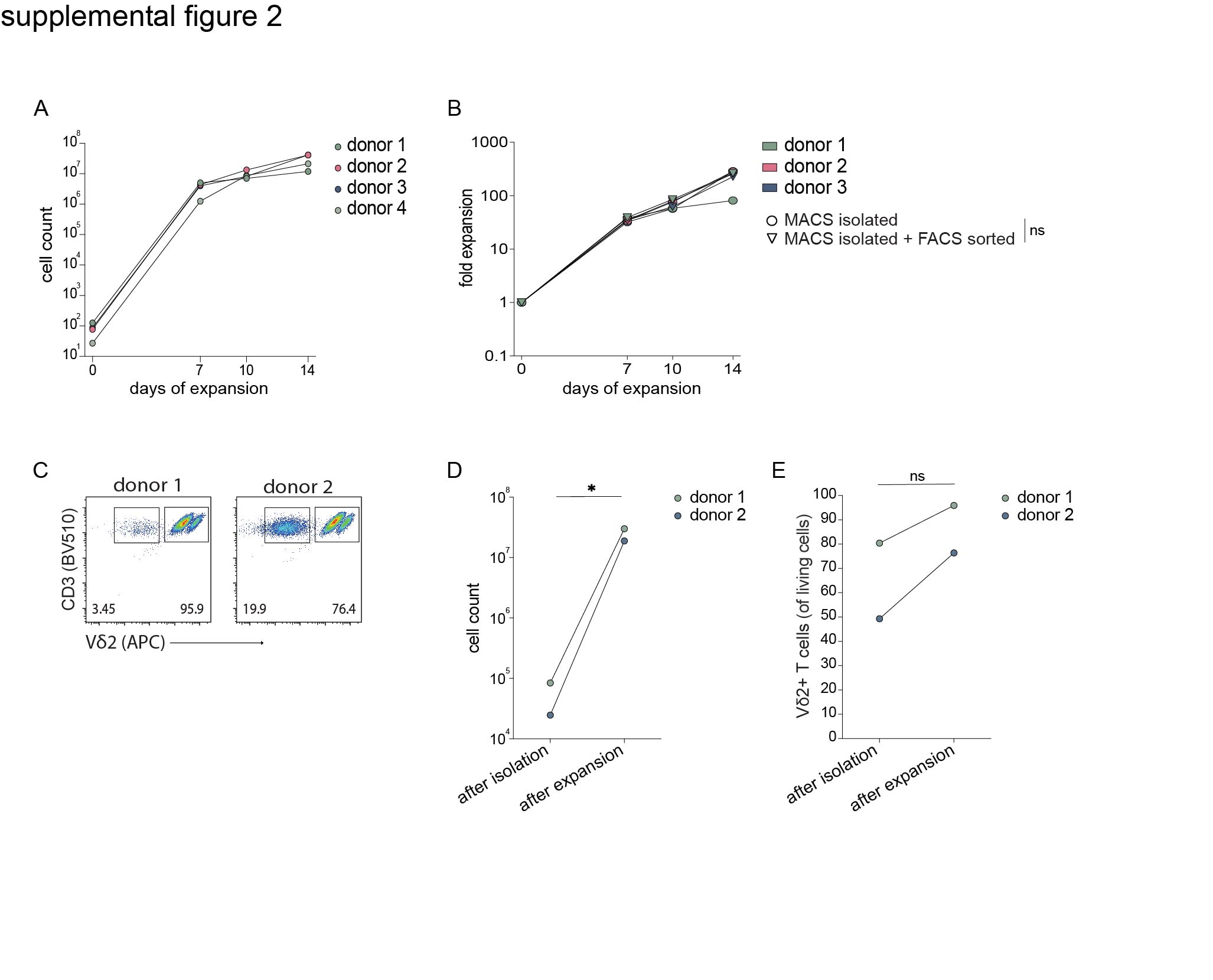
